# Transient Oxygen Exposure Causes Profound and Lasting Changes to a Benzene-Degrading Methanogenic Community

**DOI:** 10.1101/2022.04.11.487956

**Authors:** Shen Guo, Courtney R. A. Toth, Fei Luo, Xu Chen, Johnny Xiao, Elizabeth A. Edwards

**Affiliations:** Department of Chemical Engineering and Applied Chemistry, University of Toronto, 200 College Street, Toronto, Ontario M5S 3E5, Canada

**Keywords:** Benzene, anaerobic, bioremediation, methanogenesis, oxygen, *Pseudomonas*, cell decay

## Abstract

We investigated the impact of oxygen on a strictly anaerobic, methanogenic benzene-degrading enrichment culture derived decades ago from oil-contaminated sediment. The culture includes a benzene fermenter from Deltaproteobacteria Candidate clade Sva0485 (referred to as ORM2) and methanogenic archaea. A relatively small one-time injection of air, simulating a small leak into a batch culture bottle, had no measurable impact on benzene degradation rates, although retrospectively, a tiny enrichment of aerobic taxa was detected. A subsequent 100 times larger injection of air stalled methanogenesis and caused drastic perturbation of the microbial community. A benzene-degrading *Pseudomonas* became highly enriched and consumed benzene and all available oxygen. Anaerobic benzene-degrading ORM2 cell numbers plummeted during this time; re-growth and associated recovery of methanogenic benzene degradation took almost one year. These results highlight the oxygen-sensitivity of this methanogenic culture and confirm that the mechanism for anaerobic biotransformation of benzene is independent of oxygen, fundamentally different from established aerobic pathways, and is carried out by distinct microbial communities. The study further highlights the importance of including microbial decay in characterizing and modelling and mixed microbial communities.

**SYNOPSIS:** Methanogenic benzene degradation in a highly enriched anaerobic consortium was inhibited for a year after transient exposure to oxygen, causing mass decay of benzene-fermenting bacteria.

**GRAPHIC FOR ABSTRACT ART:** 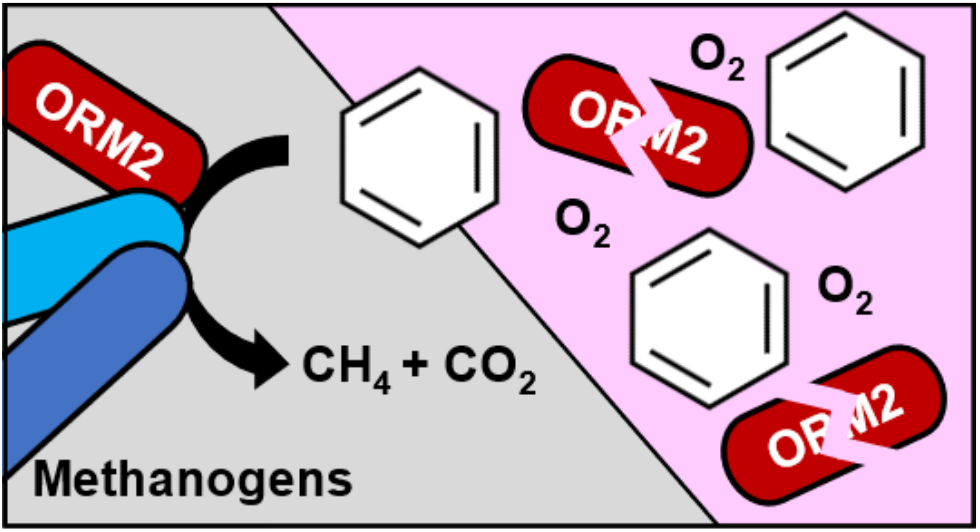

## INTRODUCTION

Benzene is a widespread and toxic pollutant notorious for its persistence in anaerobic environments including contaminated aquifers and deep sediments. Once thought to be completely recalcitrant in the absence of molecular oxygen, four decades of research now support the contrary. Anaerobic transformation of [^14^C] benzene to labelled CO_2_ was first documented in studies in the aftermath of the Amoco Cadiz oil spill in 1980 (Ward et al., 1980). Years later, this methanogenic activity was reproduced in several laboratory-scale experiments (Vogel and Grbić-Galić, 1986; Wilson et al., 1986; Grbić-Galić and Vogel, 1987; Kazumi et al., 1997; Weiner and Lovley, 1998). Oxidation of [^14^C] benzene to labelled CO_2_ was also stoichiometrically linked to the reduction of ferric iron (Coates et al., 1996; Holmes et al., 2011), nitrate (Burland and Edwards, 1999; Chakraborty et al., 2005) and sulfate (Edwards and Grbić-Galić, 1992; Lovley et al., 1995; Coates et al., 1996; Phelps et al., 1996; Kazumi et al., 1997), catalyzed by a handful of specialized microorganisms in enrichment cultures as reviewed in Vogt et al. (2011) and Toth et al. (2021). Although an enzymatic mechanism for anaerobic benzene activation has still not been established, proposed oxygen-independent transformation reactions include hydroxylation (Vogel and Grbić-Galić, 1986; Caldwell and Suflita, 2000; Zhang et al., 2013), carboxylation (Caldwell and Suflita, 2000; Kunapuli et al., 2008; Abu Laban et al., 2010; Luo et al., 2014), or activation to benzoyl-CoA involving Wood-Ljungdahl pathway intermediates (Devine, 2013). The existence of at least two different activation mechanisms is predicted from stable isotope fractionation experiments (Mancini et al., 2008).

Benzene biodegradation proceeds more readily under aerobic conditions (Atlas, 1981) and is usually preferred for bioremediation applications. In the subsurface, the caveat becomes supplying enough oxygen to drive complete benzene attenuation. Low permeability contaminated aquifers rapidly deplete dissolved O_2_, creating anaerobic zones where benzene remains recalcitrant. In situ injection of oxygen is inefficient, as O_2_ is poorly soluble in water and demand from contaminants and other reduced species is large. Other technologies such as traditional excavation or pump-and-treat strategies are costly and disruptive to the site, and not always feasible at large or deep sites. Recent studies have proposed injecting other agents such as nanoparticles (e.g., metal peroxides, zero valent iron) into the zone of contamination (Otto et al., 2008; Lu et al., 2017; Galdames et al., 2020), but these can be limited by difficulties in distribution, aggregation, maintaining activity and cost. In the end, anaerobic bioremediation may be a more economical and sustainable strategy for such locations. Decontamination can be achieved anaerobically by encouraging the growth of certain native microorganisms which can degrade contaminants (biostimulation) or by adding exogenous culture (bioaugmentation) (Reinhard et al., 1997; Cunningham et al., 2001; Major et al., 2002; Löffler and Edwards, 2006; Toth et al., 2021).

Perhaps ironically, one of the questions we are most frequently asked is if successful benzene bioremediation can truly be independent of oxygen. This skepticism is in part due the chemical stability of its unsubstituted aromatic ring and the long history of difficulties cultivating anaerobic benzene-degrading cultures (Meckenstock et al., 2016). Several reports of oxygen-generating reactions in an otherwise anoxic environment also contribute to this apprehension (Bruce et al., 1999; Weelink et al., 2008; Mehboob et al., 2009; Ettwig et al., 2010; Atashgahi et al., 2018). For example, nitrate and chlorate dismutation reactions have been shown to generate molecular oxygen internally that can then be used in benzene activation with powerful O_2_-dependent oxygenases (Weelink et al., 2008; Oosterkamp et al., 2013; Atashgahi et al., 2018). However, mono- and dioxygenase genes and associated aerobic pathways have never been detected in the metagenomes of some benzene-degrading anaerobic cultures (Abu Laban et al., 2010; Devine, 2013; Luo et al., 2014; Luo et al., 2016). Others have suggested that reactions with iron can generate radical oxygen species that may oxidize benzene (Kunapuli et al., 2008). While this can occur, it’s highly unlikely to drive benzene degradation at consistent rates and stoichiometry indefinitely, especially in longstanding enrichment cultures (Luo et al., 2014; Luo et al., 2016; Toth et al., 2021).

The objective of this study was to understand how the input of a little bit of oxygen into an active methanogenic benzene-degrading enrichment culture would affect degradation rates and impact its microbial community composition. This enrichment culture and its key benzene-degrading fermenter (ORM2, an unclassified member of the Deltaproteobacteria Candidate clade Sva0485) was recently described in detail in Toth et al. (2021). Nearly identical 16S rRNA gene sequences to ORM2 were also found in benzene-degrading methanogenic cultures from Japan (Sakai et al., 2009; Noguchi et al., 2014) and in microcosms from China (Qiao et al., 2018). From our results we determined that addition of oxygen stimulated the growth of a benzene-degrading *Pseudomonas* that was undetectable in the culture prior to oxygen exposure. Furthermore, oxygen addition resulted in dramatic reductions in ORM2 cell numbers and declines of other bacteria and archaea that are vital to this culture. Recovery of methanogenic benzene degradation took almost an entire year after anoxic conditions were re-established, attributed to the rapid decay of ORM2 cells and subsequent long doubling times to re-establish initial (pre-oxygen) concentrations. These data clearly refute any possible involvement of oxygen in benzene activation by ORM2.

## MATERIALS AND METHODS

### Methanogenic benzene-degrading culture DGG-B

DGG-B is a large-scale culture lineage derived from the lab-scale methanogenic benzene-degrading consortium OR, established over 20 years ago from site materials from an oil refinery in Oklahoma, USA (Nales et al., 1998; Toth et al., 2021). Large scale DGG-B cultures are grown at SiREM laboratories (Guelph, ON) in 100-L stainless steel vessels containing a defined anaerobic, mineral medium (Ulrich and Edwards, 2003) and benzene as their sole carbon and energy source (aqueous concentration of 25 mg/L, approximately 32 mmol per vessel) added every 4-6 weeks or as needed. The growth medium contains amorphous iron sulfide (FeS) as a reducing agent and resazurin as a redox indicator. DGG-B is currently undergoing bench-scale and field-scale testing to evaluate its potential for bioaugmentation of benzene-contaminated groundwater (Toth et al., 2021).

### Experimental setup and sampling

In May 2017, an aliquot (^~^1 L) of DGG-B was transported in a sealed bottle from SiREM labs to the University of Toronto for testing. Culture portions (30 mL) were dispensed into twenty-three 40 mL clear glass screw-cap bottles and amended with neat benzene (4.1 μmol/bottle) for an initial aqueous concentration of 10 mg/L. Bottles were sealed with Teflon Mininert screw caps and incubated in a Coy anaerobic glovebox (supplied with 10% H_2_, 10% CO_2_, and 80% N_2_) at room temperature (^~^23 °C). Benzene depletion and methane formation was monitored and bottles were reamended with benzene when concentrations dropped below 5 mg/L. After ^~^9 months of monitoring all the bottles, five bottles (designated Bottles 1 – 5) with similar and consistent benzene degradation rates (^~^0.3 μmol/bottle/day or about 0.8 mg/L/day) were selected for inclusion in this oxygen tolerance experiment.

Experimental treatments for each bottle are provided in Table 1. Day 0 of the experiment was March 19^th^, 2018, when each of the 5 similar bottles was refed and closer monitoring began. Bottles 1 to 3 were amended with an initial 0.1 mL dose of room air (^~^78% N_2_ and 21% O_2_) on Day 39 or 53 of this experiment by syringe injection. Subsequently, Bottles 1 and 2 received one (Bottle 1, Day 94) or two successive (Bottle 2, Days 94 and 224) additional doses of approximately 10 mL air (Table 1), achieved by opening (for 15 s) and closing the bottle cap to replace the headspace volume with air. As illustrated in Figure 1, the addition of 10 mL air made the resazurin in the culture medium turn pink (oxidized form). Once the resazurin returned to its colourless (reduced) form, we assumed all available O_2_ was depleted. Two bottles served as positive controls (Bottles 4 and 5) that were maintained in the glovebox as usual and never exposed to oxygen. Bottle 6 (medium only) was prepared as an anaerobic sterile control that was then exposed to 10 mL air on Day 16. Active bottles were incubated for up to 1225 days.

**Table 1.**
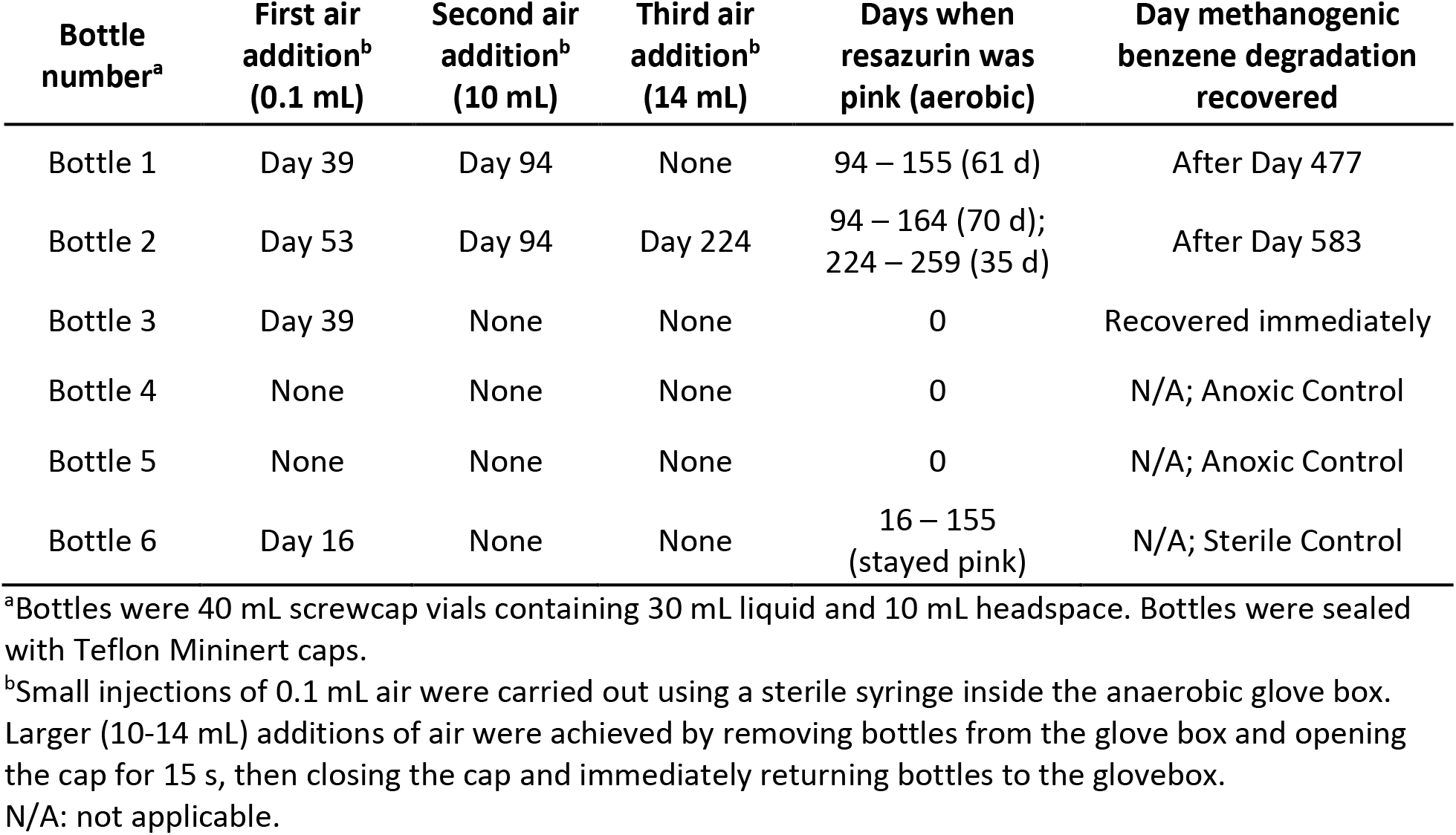
Layout of experimental treatments and overview of results.

**Figure 1.**
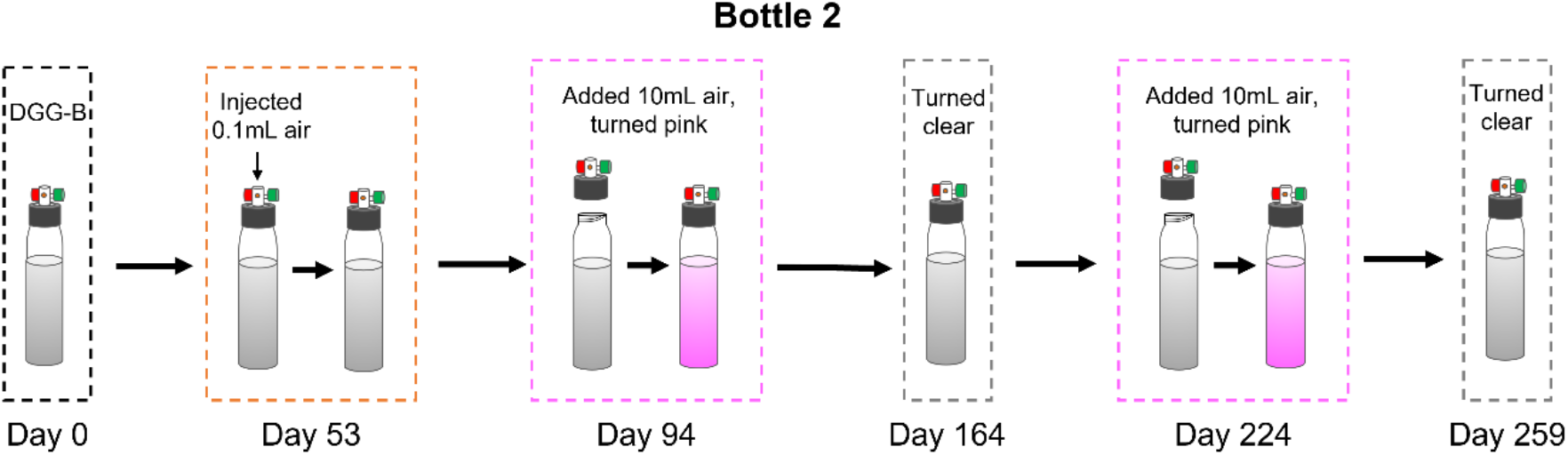
Illustration of oxygen treatment performed with DGG-B culture in Bottle 2.

### DNA extraction and molecular analyses

Bottles 1 – 5 were routinely sampled for molecular analyses. Cell pellets (10,000 × g centrifugation, 15 min) from 1 mL of culture were frozen at −80°C for later DNA extraction using the DNeasy PowerSoil Kit (Qiagen) according to the manufacturer’s procedure. Concentrations of extracted DNA were verified using a Nanodrop ND-1000 spectrophotometer (Thermo Fisher Scientific).

DNA samples were assayed by quantitative PCR (qPCR) using previously established primer pairs for total Bacteria, total Archaea, ORM2, and *Candidatus* Nealsonbacteria (previously referred to as OD1 and Parcubacteria, Table S1). Amplification of targeted genes was carried out in duplicate 20 μL reactions containing 10 μL of SsoFast™ EvaGreen^®^ Supermix (Bio-Rad Laboratories, Hercules, CA), 2 μL of DNA template, 500 nM of each designated forward and reverse primer, and UV-treated UltraPure distilled water (Thermo Fisher Scientific). Serial dilutions of plasmids containing corresponding 16S rRNA gene fragments were used to generate standard curves. qPCR reactions were performed using a CFX96 real-time thermal cycler (Bio-Rad Laboratories) using the following thermocycling conditions: an initial denaturation step at 98°C for 2 min, 40 cycles of 98°C for 5 s and T_m_ (see Table S1) for 10 s, followed by melt curve analysis (65-95°C with an increase of 0.5°C every 5 s). qPCR results were processed with CFX Manager software (Bio-Rad Laboratories).

Finally, aliquots of DNA extracts were shipped on dry ice to the Genome Quebec Innovation Centre for Illumina MiSeq 300PE (paired-end) 16S rRNA amplicon sequencing. All amplicon sequence reads were generated using modified, “staggered end” 16S rRNA gene primers 926F and 1392R (Table S1) as previously described in Toth et al. (2021). Read processing and sequencing analyses were performed in QIIME 2 version 2020.11 (Callahan et al., 2016). The SILVA SSU 132 database (Quast et al., 2013) was used to classify the resulting amplicon sequence variants (ASVs). ASVs with the same taxonomy are ranked by abundance, where ASV1 is the most abundant sequence variant recovered across all DNA samples. Two sequence variants (ASV IDs c0bc311ae7b8da92ad4208a5c83926bf and 29ca2edb9ec3825021ea0fc78298c83c) were reclassified from *Ca*. Yanofskybacteria to *Ca*. Nealsonbacteria based on matching nucleotides sequences to a complete (closed) genome recovered from the DGG-B/OR consortium (Chen et al., data in preparation; closed genome of *Ca*. Nealsonbacteria available in JGI/IMG, taxon ID 2791354853). Raw amplicon sequence reads were deposited to the National Center for Biotechnology Short Read Archive (SRA) under BioProject PRJNA807302.

### Analytical procedures

Headspace samples (300 μL) from experimental bottles were injected into a Hewlett-Packard 5890 Series II gas chromatograph equipped with GSQ 30 m × 0.53 mm I. D. PLOT column (J & W Scientific) and a flame ionization detector (GC-FID) to quantify benzene and methane as previously described (Luo et al., 2016). Benzene and methane concentrations were monitored over time in Bottles 1 – 6, from before exposure to air until after anaerobic benzene degradation resumed; all data are provided in Table S2.

### Microbial data analysis

Results of qPCR analyses of absolute abundances of targeted microbial groups (16S rRNA gene copies per mL culture) and overall community composition (from 16S rRNA gene amplicon sequencing) are summarized in Tables S3 and S4a to S4d. We also estimated the absolute abundance of all bacterial or archaeal amplicon sequence variants (ASV) detected in this study by taking the total absolute bacterial or archaeal abundance by qPCR multiplied by the relative abundance within each Domain from amplicon sequencing (Tables S5 and S6). Abundances measured using taxon-specific qPCR were similar to abundance estimated from multiplying general bacteria or archaea qPCR results by the relative abundance obtained from amplicon sequencing. Concentrations (in 16S rRNA gene copies/mL over time) for most abundant ASVs in select bottles are highlighted in main text figures (Figures 2 and 3) and for all other active bottles and major ASVs in Supplementary Figures. The amplicon sequence of one ASV of interest (*Pseudomonas* ASV1, 466 bp) was used to build a maximum likelihood tree in Geneious 8.1.9 using a GTRGAMMA model and 100 bootstrap replicates.

**Figure 2.**
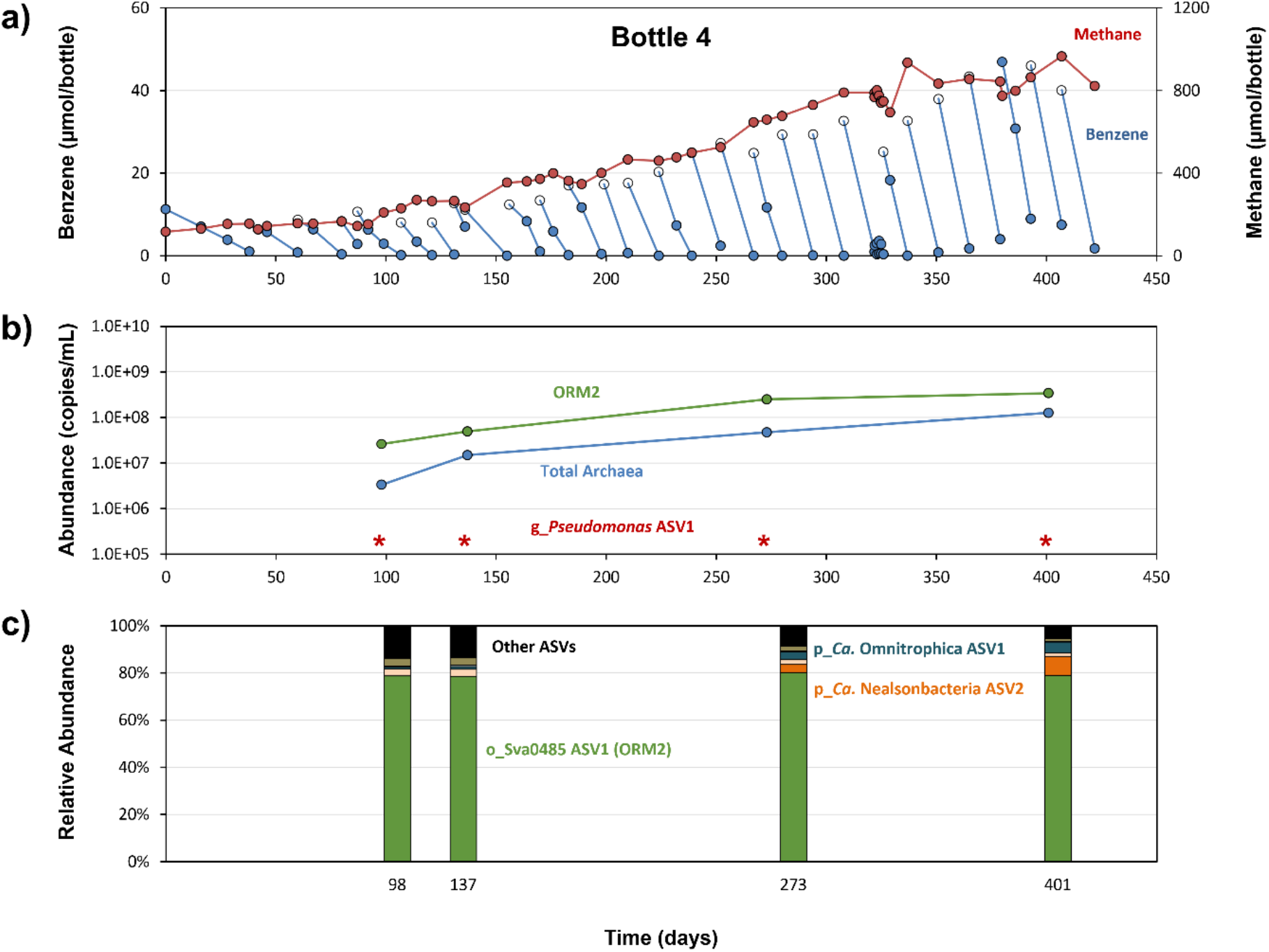
Benzene and methane concentrations and microbial profiles in anoxic control Bottle 4 over time. Panel a) presents benzene depletion (blue) and methane production (red) measured by GC-FID, where closed circles denote measured concentrations while open circles are expected benzene concentrations based on the amount fed. The center panel b) shows the absolute abundances of targeted 16S rRNA gene copies. ORM2 and Total Archaea were analyzed by qPCR; abundance of *Pseudomonas* ASV1 was calculated based on the total Bacteria qPCR abundance (Table S3) multiplied by relative abundance obtained from amplicon sequencing (Table S5). Abundances below quantifiable limits are designated by stars (*). The bottom panel c) summarizes the relative bacterial community composition measured in the bottle.

**Figure 3.**
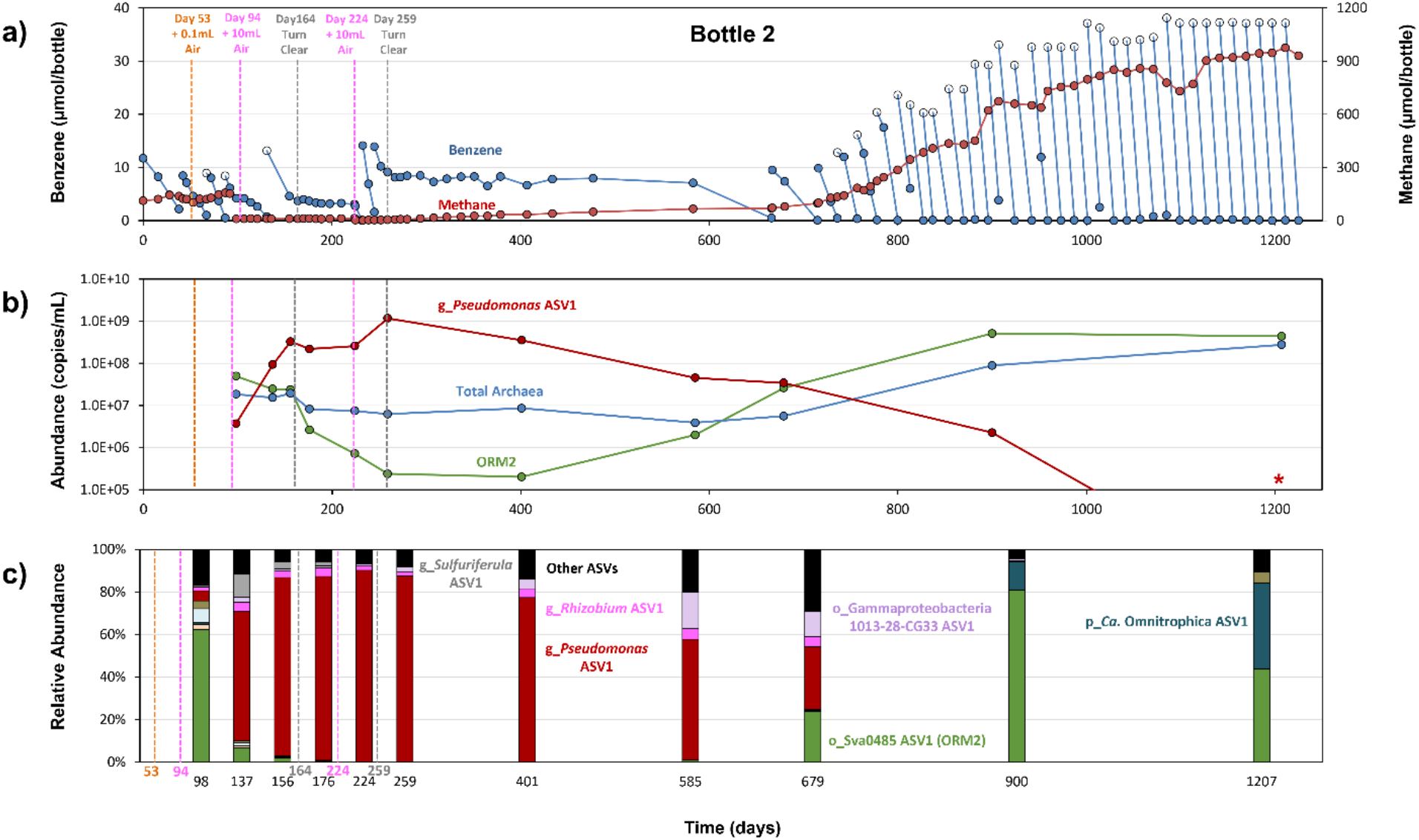
Benzene and methane concentrations and microbial profiles in oxygen-exposed Bottle 2 over time. Panel a) presents benzene depletion (blue) and methane production (red) measured by GC-FID, where closed circles denote measured concentrations while open circles are expected benzene concentrations based on the amount fed. The center panel b) shows the absolute abundances of targeted 16S rRNA gene copies. ORM2 and Total Archaea were analyzed by qPCR; abundance of *Pseudomonas* ASV1 was calculated based on the total Bacteria qPCR abundance (Table S3) multiplied by relative abundance obtained from amplicon sequencing (Table S5). Abundances below quantifiable limits are designated by stars (*). The bottom panel c) summarizes the relative bacterial community composition measured in the bottle.

## RESULTS AND DISCUSSION

### Methanogenic benzene degradation and microbial growth in bottles not exposed to oxygen

DGG-B cultures in Bottles 4 and 5 were maintained as usual in the anaerobic glovebox, without exposure to oxygen. Benzene was repeatedly degraded throughout the monitoring period, with a gradual increase in rate from 0.8 mg/L/day to 6 mg/L/day over 422 days (Figures 2a and S1a). In the first 200 days, the ratio of methane produced to benzene consumed averaged 3.5 ± 0.1 mol/mol (Table S2), nearly identical to the expected stoichiometry shown in Equation 1, where benzene (C_6_H_6_) is converted to methane (CH_4_), CO_2_ and cells (C_5_H_7_O_2_N). Similar ratios have been previously reported for consortia maintained in our laboratory (Ulrich and Edwards, 2003; Luo et al., 2016) and elsewhere (Kazumi et al., 1997; Sakai et al., 2009). After 200 days, measured methane concentrations began to plateau and were underestimated due to pressure buildup and associated losses from sampling.

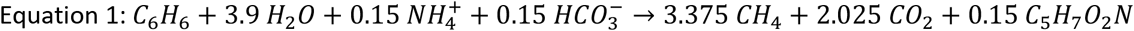

Dominant microbial phylotypes in the DGG-B/OR consortium include Deltaproteobacteria Candidate clade Sva0485 (ORM2), acetoclastic *Methanosaeta* and hydrogenotrophic *Methanoregula* that are required for complete biodegradation of benzene to CH_4_ and CO_2_ (Devine, 2013; Luo et al., 2016; Toth et al., 2021). A member of the Candidate Phyla Radiation (*Ca*. Nealsonbacteria, previously referred to as OD1 and Parcubacteria) has also been consistently found in the culture (Luo et al., 2016; Toth et al., 2021) and is thought to be involved in biomass recycling (Chen et al., data in preparation). Absolute abundance was tracked by qPCR and is shown in Figure 2b (Bottle 4) and Figure S1b (Bottle 5). Turning next to 16S rRNA gene amplicon sequencing, three ASVs increased the most; Sva0485 Deltaproteobacteria ASV1 (ORM2), *Methanosaeta* ASV1, and *Methanoregula* ASV1 (Figures 2c and S1c; Tables S5 and S6). Other high abundance ASVs belonged to *Ca*. Nealsonbacteria and *Ca*. Omnitrophica (previously referred to as OP3), a candidate phylum that may also implicated in biomass recycling (Suominen et al., 2021)(Figures 2c and S1c; Tables S3 and S5). Overall, the observed benzene degradation rates and growth of ORM2, methanogens and candidate phyla were as expected in these positive control bottles.

### Benzene biodegradation following oxygen exposure

In experimental bottles, the impacts of exposure to oxygen are illustrated in Figure 3a (Bottle 2), Figures S2a and S3a (Bottles 1 and 3), and summarized in Table 1. After the first addition of 0.1 mL air, the culture medium remained clear, indicating that the resazurin remained reduced. There was also no observable impact on benzene degradation rate or methane production, as seen by the data from Bottle 3 (Figure S3). We surmised that the FeS used to reduce the culture medium (15 μmol/bottle) scavenged the added O_2_ (^~^0.9 μmol/bottle) and that exceeding the oxygen demand from FeS (34 μmol/bottle) would be necessary to elicit a change in benzene degradation activity (calculations shown in Table S7). The headspace volume of the bottles was 10 mL. We reasoned that if we simply opened the bottle cap for a few seconds and then closed it, we would effectively replace the headspace volume with air. Using this approach, we added ^~^10 mL of air (^~^86 μmol/bottle of O_2_) once to Bottle 1, twice to Bottle 2, and once to medium-only (sterile) Bottle 6 (Table 1). Expectedly, the culture medium immediately turned pink and remained pink for up to 70 days in live replicates (Figures 3a, S2a, and S4). No methane production was observed when the bottles were still pink. Benzene degradation was able to proceed in Bottles 1 and 2 following a brief lag period (^~^1 week) but stopped after 61 – 70 days, coinciding with oxygen depletion (when the resazurin turned clear again). After adding a second 10 mL of air to Bottle 2, benzene degradation resumed for 35 days until O_2_ was depleted once again (Figure 3a). No benzene loss was detected in Bottle 6 after oxygen was added (Figure S4), indicating that the benzene losses seen in Bottles 1 and 2 were not the result of contamination from added air. A moles balance comparing benzene degraded during the aerobic “pink” period to the amount of available oxygen in each bottle confirmed that observed benzene depletion after adding air to the bottles was coupled to oxygen consumption (Table S7).

We continued to monitor these bottles to determine if and when anaerobic benzene degradation would re-establish. Methane production began slowly over several months, and finally methanogenic benzene degradation was measurable again by Day 583 in Bottle 1 and by Day 666 in Bottle 2 (Figures 3a and S2a; Table 1). This lag of 323-325 days was much longer than expected, even considering the known, slow doubling time of ^~^30 days for ORM2 (Ulrich and Edwards, 2003; Luo et al., 2016; Toth et al., 2021). This prompted us to take a closer look at the microbes in each bottle over time.

### Appearance of *Pseudomonas* upon exposure to oxygen

Our amplicon sequencing data immediately revealed that an ASV affiliated with *Pseudomonas* became highly enriched in Bottles 1 and 2 after adding 10 mL of air (Figures 3b, 3c, S2b, and S2c). Using Bottle 2 as an example, we can see concentrations of *Pseudomonas* ASV1 copies/mL increased rapidly in the presence of oxygen (up to 1.2 × 10^9^ copies/mL) but decayed very slowly under anoxic conditions (Figure 3b). It took until Day 1207 for concentrations of *Pseudomonas* ASV1 to fall below detectable limits. Looking closely at Bottle 3, we can see transient, low abundances of *Pseudomonas* ASV1 appear after the first 0.1 mL injection of air, even though no perturbation of the culture was observed (Figure S3b). *Pseudomonas* have not previously been detected in the DGG-B/OR consortium (Ulrich and Edwards, 2003; Luo et al., 2016; Toth et al., 2021) and no ASVs associated with *Pseudomonas* were ever detected in anoxic controls Bottles 4 and 5 (Table S4a). This ASV was likely not a contaminant from the air or growth medium used because benzene losses were never detected in medium-only control Bottle 6 (Table 1; Figure S4). *Pseudomonas* ASV1 is closely related (100% sequence identity) to several *Pseudomonas stutzeri* isolates (Figure S5), although a longer 16S rRNA gene sequence (>466 base pairs) or genome is needed to verify taxonomy. *P. stutzeri* species are metabolically versatile and can use oxygen or nitrate as a terminal electron acceptor (Lalucat et al., 2006). A handful of strains are also capable of mineralizing aromatic hydrocarbons (Grimberg et al., 1996; Ortega-Calvo et al., 2003; Heinaru et al., 2016). Considering the above analysis, benzene-degrading *Pseudomonas* likely persist below detection in the DGG-B culture and were responsible for the aerobic benzene metabolism seen in Bottles 1 and 2 (Figures 3a and S2a).

### Impact of oxygen on ORM2 and methanogens

The community composition and absolute abundance of ORM2, Archaea and Bacteria tracked in anoxic controls Bottles 4 and 5 provide a reference against which oxygen-impacted Bottles 1 – 3 were compared(Figures 2, 3, S1-S3). Prior to oxygen addition, concentrations of ORM2 in Bottle 3 (^~^9.5 × 10^7^ copies/mL) were comparable to those measured in anoxic control Bottles 4 and 5 (Table S3). After adding 10 mL of air to Bottles 1 and 2, concentrations of ORM2 steeply decreased by up to 2 orders of magnitude (Figures 3b and S2b). The addition of only 0.1 mL of air in Bottle 3 may even have resulted in a small (^~^53%) decrease in concentrations of ORM2 (Table S3). We were surprised to see that ORM2 concentrations continued to decline in Bottles 1 and 2 long after the bottles were no longer pink and presumed to be devoid of O_2_. The lowest concentrations of ORM2 measured were 2.9 × 10^6^ copies/mL on Day 401 in Bottle 1 (Figure S2b) and 2.0 × 10^5^ copies/mL on Day 401 in Bottle 2 (Figures 3b). Both concentrations are below estimated threshold concentrations of ORM2 (4.3 × 10^6^ ORM2 copies/mL) needed for measurable benzene loss in batch cultures of DGG-B (Toth et al., 2021). Concentrations of total Archaea also declined up to 1 order of magnitude between Days 98 – 585 (Figures 3b and S2b). While ORM2 abundance decreased, the abundance of total Bacteria increased by 56-80% immediately after adding 10 mL of air (Tables S3 and S5), driven by the growth of aforementioned *Pseudomonas*. After the medium in Bottles 1 and 2 turned clear, *Pseudomonas* copies began to decrease yet it still took a further 140-240 days to see net increases in ORM2 copies. This prompted us to take a closer look at observed growth and decay rates. Transient oxygen exposure had a prolonged negative impact on the microbial community structure of DGG-B.

### Estimates of cell yields, growth and decay rates for *Pseudomonas* and ORM2

We performed mass and electron balances in each bottle to compare the ratios of donor (benzene) consumption to oxygen reduction (in aerobic phases) and to methane formation (in anoxic phases), including biomass formation. Cell yields were estimated from qPCR (16S rRNA gene copies/mL), cell size (g/copy), and benzene consumption data over each time interval; these calculations are detailed in Table S8. Although qPCR can overestimate cell copy numbers by capturing DNA from dead cells (Kralik and Ricchi, 2017), we assumed this risk to be minimal since cell decay on the orders of magnitude were observed in this study. Measured values were compared to predictions shown in Equation 1 (methanogenic case) and in Equation 2 (aerobic case). In Equation 2, the fraction of electrons from benzene going to cell synthesis was 50% which is typical for aerobic processes (Rittmann and McCarty, 2001).

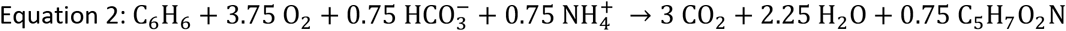

The average measured yield for *Pseudomonas* (^~^1.1 g cells per g of benzene) during the aerobic phase was remarkably close the predicted yield (Table 2). In contrast, the total yield of ORM2 and total Archaea under methanogenic conditions (^~^0.04 g cells/g benzene) was found to be well below predicted growth yield (0.22 g/g) shown in Equation 1 and in Table 2. Low growth yields have been previously reported for this culture (Luo et al., 2016; Toth et al., 2021) and are consistent with the difficulties commonly reported in trying to sustain high rates and cell concentrations in anaerobic benzene-degrading cultures.

**Table 2.**
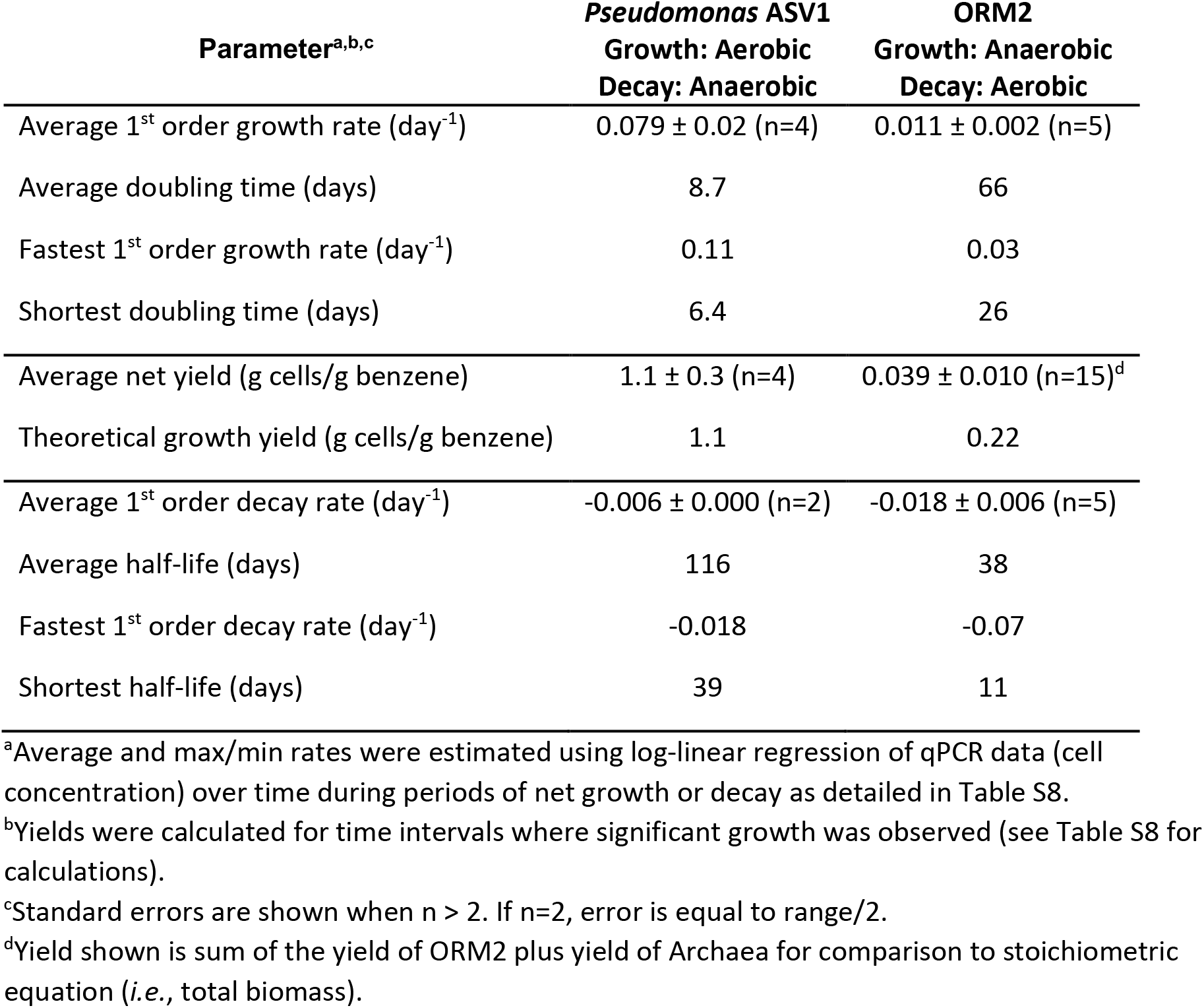
Estimated first order growth and decay rates for benzene-degrading microorganisms in DGG-B.

To explore growth and decay rates more closely, we estimated net (*i.e*., measured) cell growth and decay rates for the two benzene degraders, *Pseudomonas* and ORM2 (Table S8), assuming first order processes as an approximation. Using the slope from first order regression of microbial abundance (from qPCR) vs time over specific intervals (aerobic, anaerobic), we were able to get some estimates that reflect observations in Bottles 1 – 5 (Table S8). In Table 2, we summarize the average and fastest rates estimated and provide calculated, representative doubling times or half-lives to make comparisons easier.

Growth rates for *Pseudomonas* were substantially faster than for ORM2, best seen in Table 2 by the difference in minimum/average doubling times (6/9 days vs 26/66 days, respectively). These data are consistent with literature and previous reports for ^~^30 day doubling times for ORM2 (Ulrich and Edwards, 2003; Luo et al., 2016; Toth et al., 2021). As for observed decay rates, we see striking numbers for ORM2, particularly during aerobic phases where the fastest decay rates and shortest half-lives of only 11 days were reported (Table 2, Table S8). This highlights a substantial loss of ORM2 biomass upon exposure to oxygen. The average half-life of ORM2 (calculated during aerobic and anaerobic periods of decay) was about 38 days, which is very short and incredibly close to the organism’s known doubling time. Therefore it is not surprising that it took almost a year for the culture to recover. In previous experiments by Luo et al. (2016), decay was essentially zero for ORM2 maintained anaerobically and without benzene. ORM2 is thus highly sensitive to oxygen and reinforces the existence of a novel, still unknown pathway independent of O_2_ in methanogenic benzene degradation.

### Impact of oxygen on other microbial community members

After realizing just how sensitive ORM2 is to oxygen, we were curious how other members of the DGG-B community responded to air addition events. Putative microbes represented by ASVs were divided into Bacteria and Archaea and further categorized into one of three groups; i) ASVs that became enriched aerobically, ii) ASVs that declined in response to O_2_ exposure, and iii) ASVs that were largely unaffected (Figures S6a – S6d and S7a – S7d).

Group 1 Bacteria that were positively impacted by oxygen include the *Pseudomonas* ASV1 discussed above, as well as *Sulfurferula* ASV1, *Thiomonas* ASV1, and *Rhizobium* ASV1 (Figures S6aS6b and S6c). None of these ASVs were detectable in anoxic control Bottles 4 and 5 at any time point (Figure S6d). Members of these genera are aerobic but can survive under anoxic conditions (Vela et al., 2002; Lalucat et al., 2006; Watanabe et al., 2015; 2016). *Sulfuriferula* and *Thiomonas* are chemolithoautotrophs that oxidize some sulfur species (S^0^, thiosulfate, sulfide, and tetrathionate) with O_2_ (Watanabe et al., 2015; 2016). The FeS reducing agent in our growth medium could logically have supported the growth of *Sulfuriferula* ASV1 or *Thiomonas* ASV1 in the presence of O_2_. In our electron balance, we assumed the oxidation of FeS was abiotic but these data suggest it may instead have been biotic. Select rhizobial species can metabolize substituted aromatics including phenol, a hydroxylated form of benzene that can be produced abiotically by autooxidation (Vela et al., 2002; Kunapuli et al., 2008).

Group 2 microbes that were negatively impacted by oxygen include those we have already discussed: *Deltaproteobacteria* ASV1 (ORM2), two *Ca*. Nealsonbacteria ASVs (OD1) and *Ca*. Omnitrophica ASV1 (Figures S6a and S6b). High abundance ASVs associated with *Methanosaeta* and *Methanoregula* also decreased 1-2 orders of magnitude follow the addition of 10 mL air (Figures S7a and S7b). Concentrations of most of these bacterial and archaeal ASVs did not begin to recover until after ORM2 growth resumed and methanogenic benzene degradation was re-established. Notably, a few Group 2 ASVs in Bottles 1 and 2 never fully recovered from O_2_ exposure, including *Ca*. Nealsonbacteria ASV1 (Figures S6a and S6b), *Methanoregula* ASV2 and *Methanosaeta* ASV4 (Figures S7a and S7b), pointing to likely permanent community impacts.

Group 3 comprises largely unimpacted microbes that were in low abundance and include two unclassified *Spirochaetaceae* ASVs, one ASV belonging to an unclassified *Anaerolineaceae* genus, and one ASV belonging to hydrogenotrophic *Methanobacterium* (Figures S6a – S6d and S7a – S7d). *Spirochaetaceae* and *Anaerolineaceae* are commonly detected in anaerobic and methanogenic hydrocarbon-degrading enrichment cultures, but their roles often undefined. Some Spirochaetes are believed to mediate recycling of biomass from dead cells (Dong et al., 2018). *Anaerolineceae* are often closely associated with methanogens and may catalyze the degradation of other hydrocarbons (Liang et al., 2015; Mohamad Shahimin and Siddique, 2017). Our results suggest that the roles of Group 3 microorganisms are independent of benzene degradation.

### Implications for the field

This study has made it abundantly clear that ORM2, the chief anaerobic benzene degrader in the DGG-B/OR consortium, is strictly anaerobic and oxygen-sensitive. Transient oxygen exposure contributed to extensive, prolonged decay of ORM2 and associated Group 2 methanogens and putative biomass-recycling bacteria. A complete (closed) genome for ORM2 has been sequenced and is under investigation (JGI/IMG taxon ID 2795385393) but its biochemical mechanism for benzene transformation is still not known. Nevertheless, we can say for certain is that oxygen is not involved in anaerobic benzene degradation by this culture and probably not in other consortia harboring ORM2-like benzene degraders (Sakai et al., 2009; Noguchi et al., 2014; Qiao et al., 2018).

For years our laboratory has wondered why anaerobic benzene degraders such as ORM2 rarely proliferate in subsurface benzene-contaminated environments. We recently explored environmental co-contaminants (*i.e*., toluene and xylenes) and microbial co-dependencies as possibilities constraining their growth (Toth et al., 2021) but acknowledged that a multitude of other factors were likely involved. This study implicates transient oxygen exposure as another possibility. In the subsurface, dissolved oxygen may be introduced to anoxic zones from groundwater recharge, fluctuations in groundwater elevation, rain events, and diffusion from the vadose zone. Shallow unconfined aquifers are much more likely to see oxygen incursions. However, unlike our culture bottles, subsurface environments are highly heterogeneous and filled with microenvironments that would not see bulk oxygen concentrations. Nevertheless, careful attention to avoiding changes in redox potential appears warranted if sustained anaerobic benzene biotransformation is desired.

This study has also highlighted pronounced cell death/decay as another important factor constraining net microbial growth. Cell decay is often ignored in mathematical models, particularly in contaminant fate and transport models. It is frequently assumed that because anaerobic bacteria do not grow quickly, their decay rates must also be very low and can therefore be neglected. A rule of thumb is that decay rates are 1/10^th^ of growth rates (Rittmann and McCarty, 2001). Contrarily, we observed for ORM2 that growth and decay rates were of similar magnitude, even when oxygen was no longer present. This result is consistent with previously described challenges in achieving high cell densities and consistent benzene degradation rates in lab cultures. They could also help explain why organisms putatively associated with biomass recycling, including *Ca*. Nealsonbacteria (OD1) and *Ca*. Omnitrophica (OP3), among others, are periodically observed in high abundances in the DGG-B/OR consortium (Luo et al., 2016; Toth et al., 2021) and other anaerobic benzene-degrading consortia (Taubert et al., 2012; Melkonian et al., 2021). What factors could explain such relatively high decay rates? Possible explanations include viral and bacterial predation or microbial competition. Another explanation could be the formation of toxic reactive intermediates during benzene activation resulting in cell death, or the energy penalty afforded by this difficult reaction. Understanding the underlying mechanisms of death and decay will be key to establishing more robust cultures for bioremediation. Interestingly, comparable methanogenic enrichment cultures that degrade toluene rather than benzene do not exhibit the same struggles and thus the most likely hypothesis is that a step in the biochemical activation of benzene contributes to higher rates of decay.

## AUTHOR CONTRIBUTIONS

All authors made research and substantial intellectual contributions to the completion of this study. E.A.E., S.G. and F.L. conceived the project. S.G. established all experimental bottles and performed all GC measurements, DNA extractions and qPCR assays, with assistance from X.C. and J.X. Amplicon sequencing data processing and analyses were completed by S.G. and C.R.A.T. Cell yields, growth and decay rate estimates were completed by C.R.A.T., S.G., and E.A.E. The manuscript was written by S.G., C.R.A.T. and E.A.E. All authors contributed to manuscript revision and have approved the submitted version.

## SUPPORTING INFORMATION

Figures showing benzene and methane concentrations and microbial profiles in Bottle 5 (positive control), Bottle 1 (oxygen-amended), Bottle 3 (oxygen-amended) and Bottle 6 (sterile) over time; maximum likelihood consensus tree for *Pseudomonas;* absolute abundance of specific bacterial and archaeal groups in all bottles (PDF)

Primers used for amplification; benzene and methane concentrations; qPCR raw data including calibration curves; all gDNA samples sequenced, including ASVs and their taxonomic assignment; calculation of absolute abundances; oxygen demand stoichiometry; estimation of yields, growth and decay rates (XLSX)

## ACKNOWLEDGEMENTS

This study was funded by Genomic Application Partnership Program (GAPP) grants awarded to E.A.E. (Project IDs OGI-102 and OGI-173), which were supported by Genome Canada, Ontario Genomics, the Government of Ontario, Mitacs Canada, SiREM, Alberta Innovates, Federated Co-operatives Limited, and Imperial Oil. The authors would like to thank SiREM (Guelph, Ontario N1G 3Z2, Canada) for supplying DGG-B culture for oxygen tolerance testing. We would also like to acknowledge the laboratories of Dr. Neil Thomson (University of Waterloo), Dr. Ania Ulrich (University of Alberta), SiREM, and Innotech Alberta (Edmonton, Alberta T6N 1E4, Canada) for providing feedback and intellectual contributions to this study during weekly GAPP consortium meetings.

