## Supplemental Information for "Transient Oxygen Exposure Causes Profound and Lasting Changes to a Benzene-Degrading Methanogenic Community"

Number of pages: 13

Number of Supplementary Tables: 8

*Supplementary Tables (S1-S8) are in separate Excel file

Number of Supplementary Figures: 7 (see below)

### TABLE OF CONTENTS


**Supplementary Tables (all in accompanying Excel file)**

**Table S1:** Primers used for quantification of targeted 16S rRNA genes.

**Table S2:** Summary of benzene and methane concentrations (mg/L) and their calculated (µmol) in experimental bottles.

**Table S3:** Concentrations of target microorganisms in experimental bottles as measured by qPCR of 16S rRNA genes.

**Table S4:** List of all gDNA samples amplified by Illumina sequencing in this study;
- **4a**: raw reads (read numbers) for all ASVs and their taxonomic assignment.
- **4b**: percentage of ASVs and their taxonomic assignment.
- **4c**: bacterial ASVs as a percent of all bacterial ASVs.
- **4d:** archaeal ASVs as a percent of all archaeal ASVs.

**Table S5:** List of absolute abundance of each bacterial ASV.

**Table S6:** List of absolute abundance of each archaeal ASV.

**Table S7:** Stoichiometric balances for oxygen in experimental bottles.

**Table S8**: Estimation of ORM2, *Pseudomonas* and Archaea yields, growth and decay rates in experimental bottles.

**Supplementary Figures**

**Figure S1:** Benzene and methane concentrations and microbial profiles in positive control Bottle 5 over time.

**Figure S2:** Benzene and methane concentrations and microbial profiles in Bottle 1 over time.

**Figure S3:** Benzene and methane concentrations and microbial profiles in Bottle 3 over time.

**Figure S4:** Benzene and methane concentration profiles in sterile control Bottle 6.

**Figure S5:** Maximum likelihood consensus tree showing the affiliation of *Pseudomonas* ASV1 to reference *Pseudomonas* isolates.

**Figure S6:** Absolute abundance of specific bacterial groups;
- **6a**: in Bottle 1.
- **6b**: in Bottle 2.
- **6c**: in Bottle 3.
- **6d**: in Bottles 4 and 5.

**Figure S7**: Absolute abundance of specific archaeal groups;
- **7a**: in Bottle 1.
- **7b**: in Bottle 2.
- **7c**: in Bottle 3.
- **7d**: in Bottles 4 and 5.

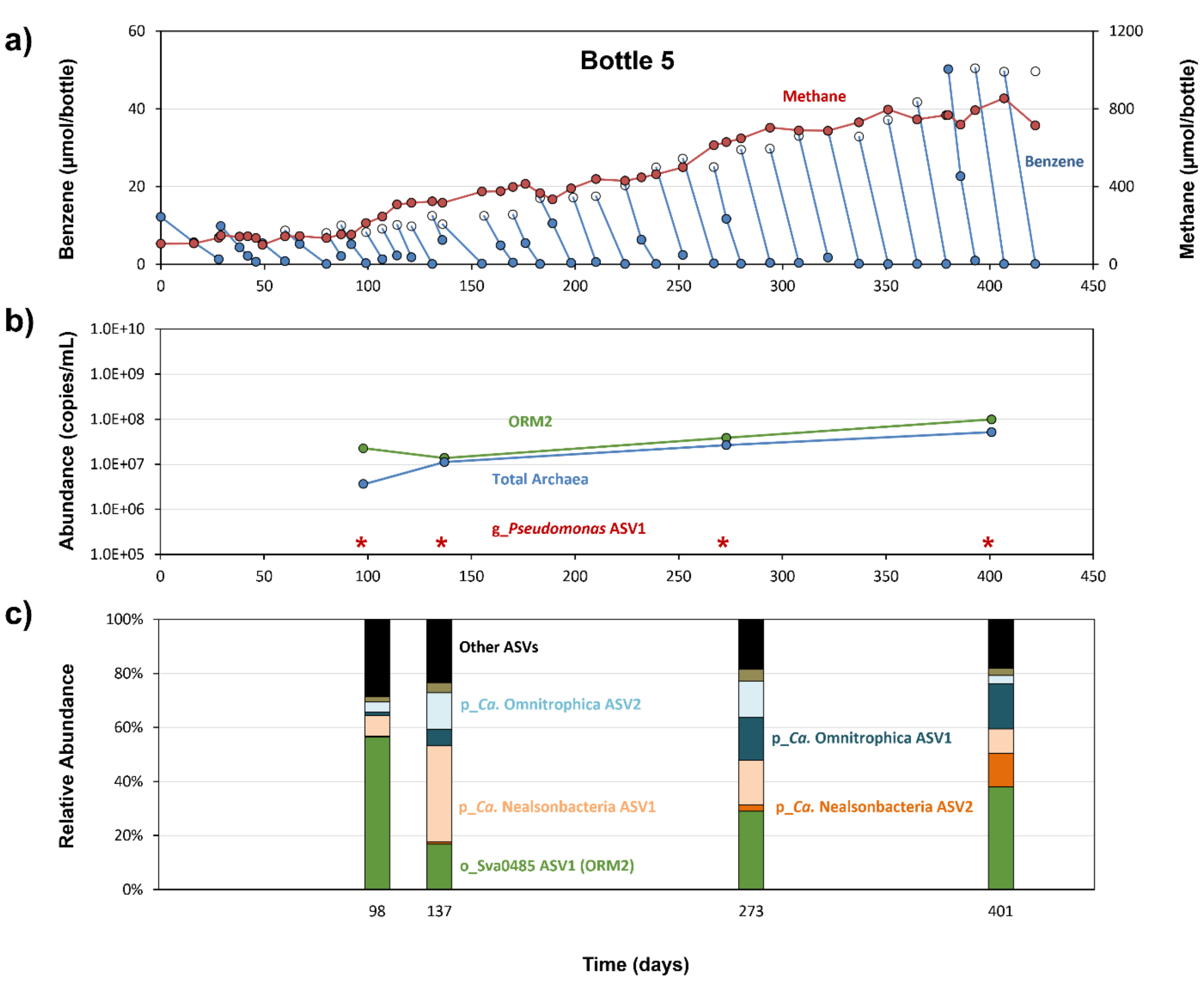


**Figure S1**. Benzene and methane concentrations and microbial profiles in anoxic control Bottle 5 over time. Panel a) presents benzene depletion (blue) and methane production (red) measured by GC-FID, where closed circles denote measured concentrations while open circles are expected benzene concentrations based on the amount fed. The center panel b) shows the absolute abundances of targeted 16S rRNA gene copies. ORM2 and Total Archaea were analyzed by qPCR; abundance of *Pseudomonas* ASV1 was calculated based on the total Bacteria qPCR abundance (Table S3) multiplied by relative abundance obtained from amplicon sequencing (Table S5). Abundances below quantifiable limits are designated by stars (*). The bottom panel c) summarizes the relative bacterial community composition measured in the bottle.

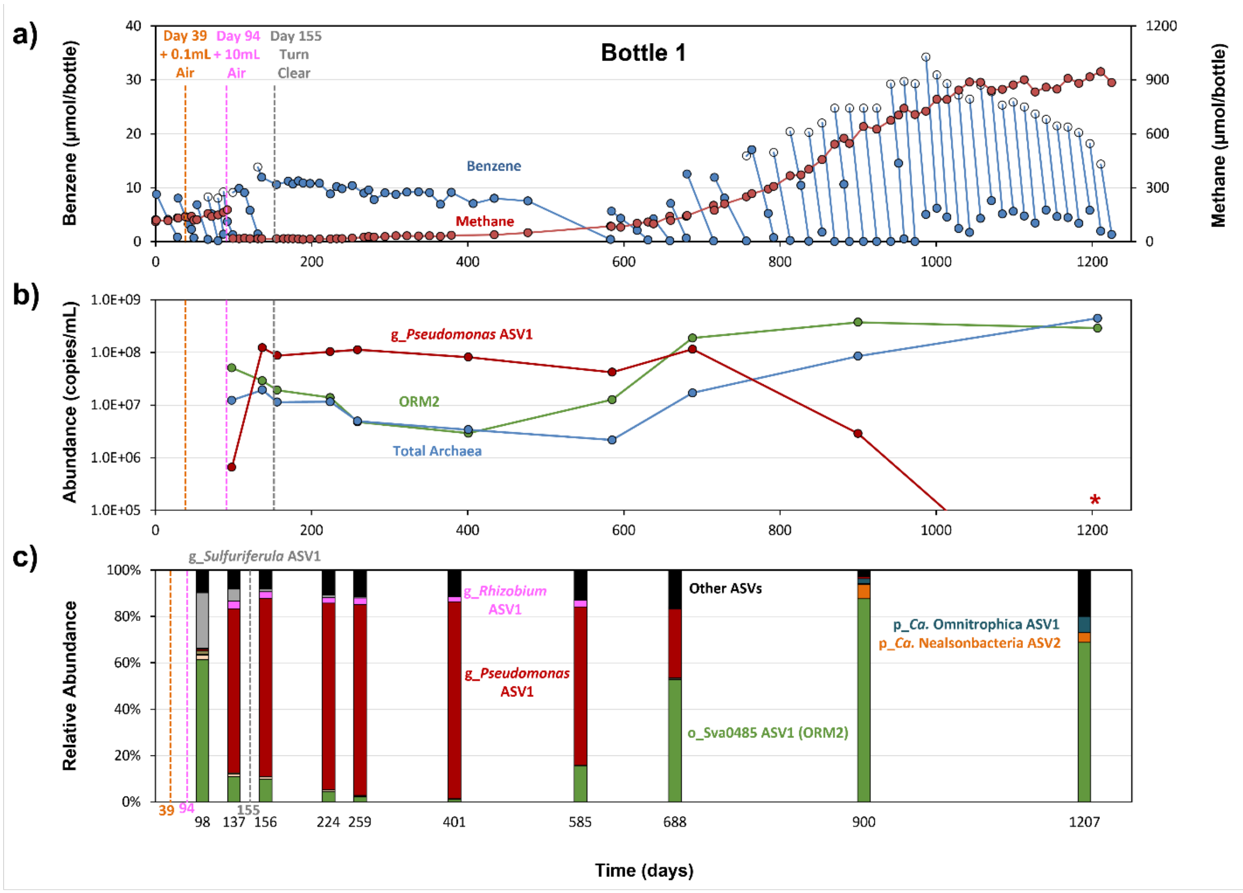


**Figure S2**. Benzene and methane concentrations and microbial profiles in oxygen-exposed Bottle 1 over time. Panel a) presents benzene depletion (blue) and methane production (red) measured by GC-FID, where closed circles denote measured concentrations while open circles are expected benzene concentrations based on the amount fed. The center panel b) shows the absolute abundances of targeted 16S rRNA gene copies. ORM2 and Total Archaea were analyzed by qPCR; abundance of *Pseudomonas* ASV1 was calculated based on the total Bacteria qPCR abundance (Table S3) multiplied by relative abundance obtained from amplicon sequencing (Table S5). Abundances below quantifiable limits are designated by stars (*). The bottom panel c) summarizes the relative bacterial community composition measured in the bottle.

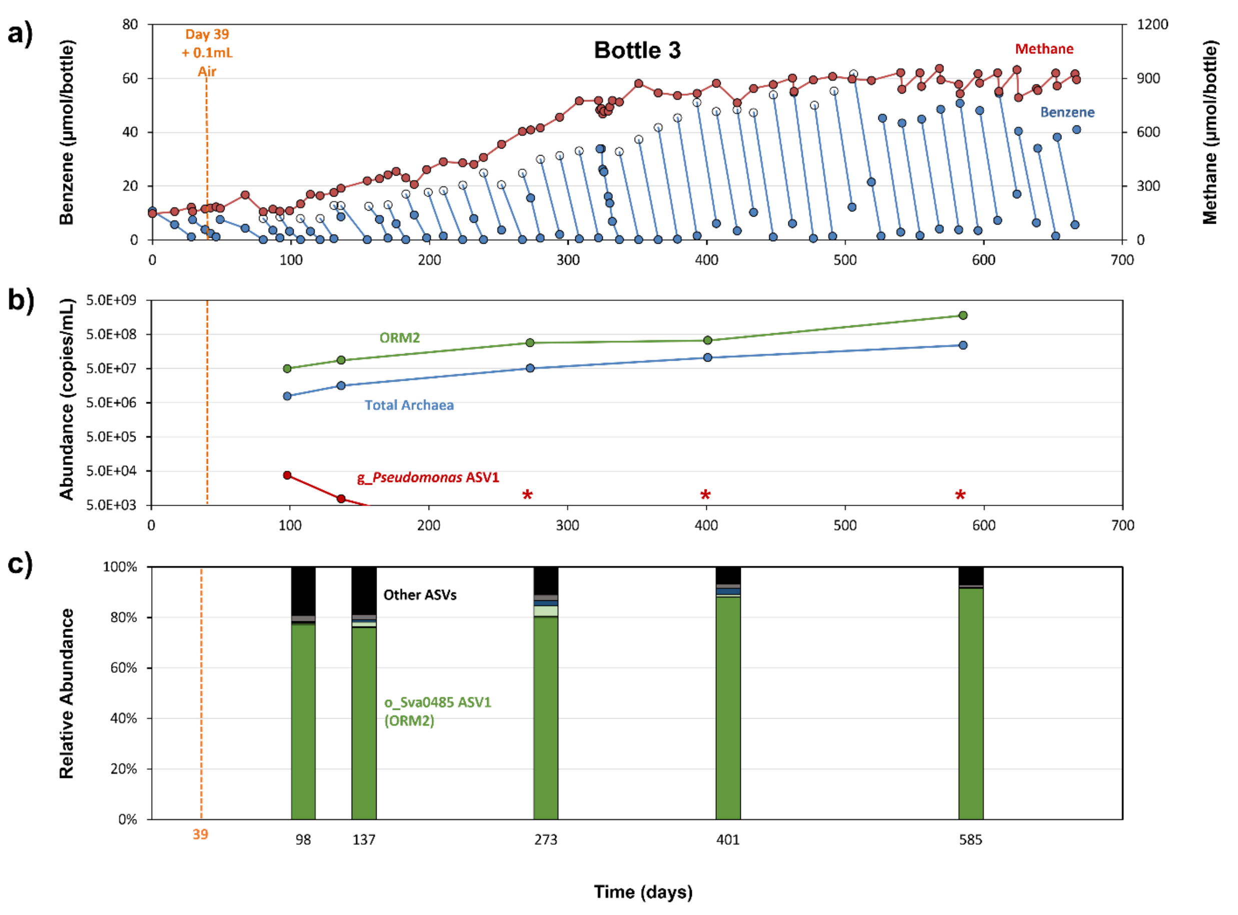


**Figure S3**. Benzene and methane concentrations and microbial profiles in Bottle 3 over time. Panel a) presents benzene depletion (blue) and methane production (red) measured by GC-FID, where closed circles denote measured concentrations while open circles are expected benzene concentrations based on the amount fed. The center panel b) shows the absolute abundances of targeted 16S rRNA gene copies. ORM2 and Total Archaea were analyzed by qPCR; abundance of *Pseudomonas* ASV1 was calculated based on the total Bacteria qPCR abundance (Table S3) multiplied by relative abundance obtained from amplicon sequencing (Table S5). Abundances below quantifiable limits are designated by stars (*). The bottom panel c) summarizes the relative bacterial community composition measured in the bottle.

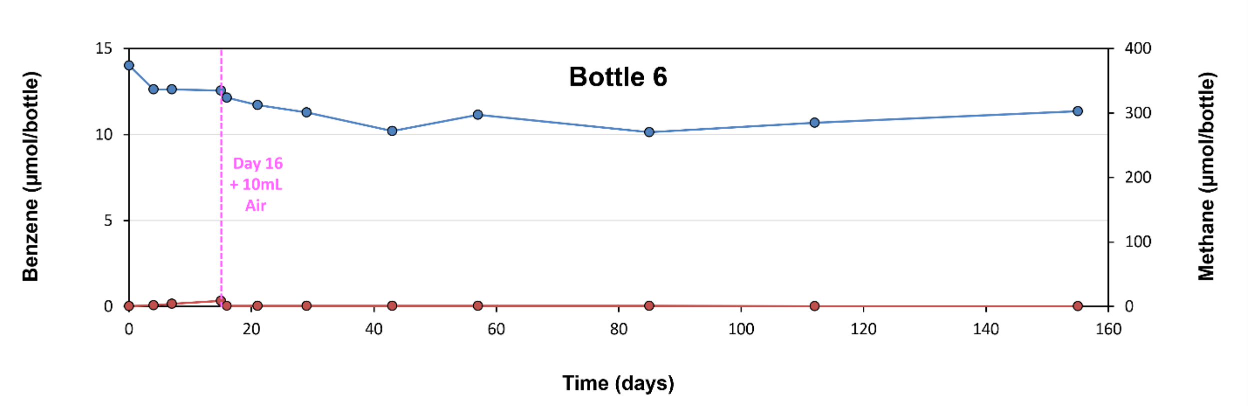


**Figure S4**. Benzene and methane concentration profiles in sterile control Bottle 6. Closed circles denote concentrations of benzene (blue) and methane (red) measured by GC-FID. As no benzene losses or methane production was observed, no molecular samples were collected.

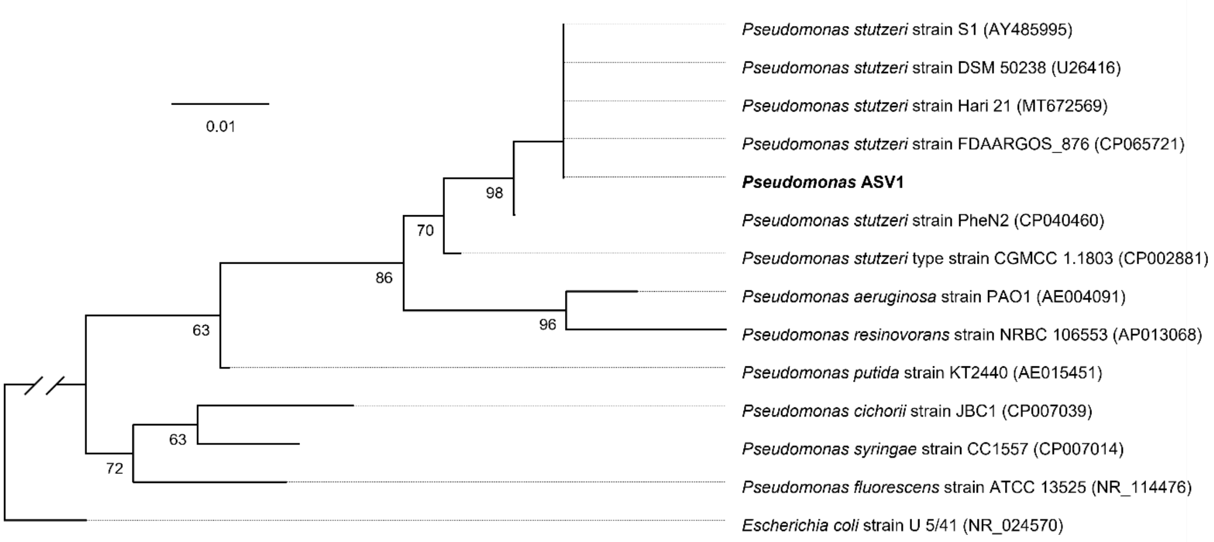


**Figure S5**: Maximum likelihood (ML) consensus tree showing the affiliation of *Pseudomonas* ASV1 (bold) to reference *Pseudomonas* isolates. Full-length and partial 16S rRNA gene sequences were aligned using MUSCLE (Edgar, 2004) and the ML tree was constructed from the resulting alignment in RAxML version 7.2.8 (Stamatakis, 2014) in Geneious version 8 using a GTRGAMMA model and 100 bootstrap replicates. Bootstraps <60% are not shown.

**Figure 6 comprises a set 4 related figures (a,b,c,d) each for different bottles.**

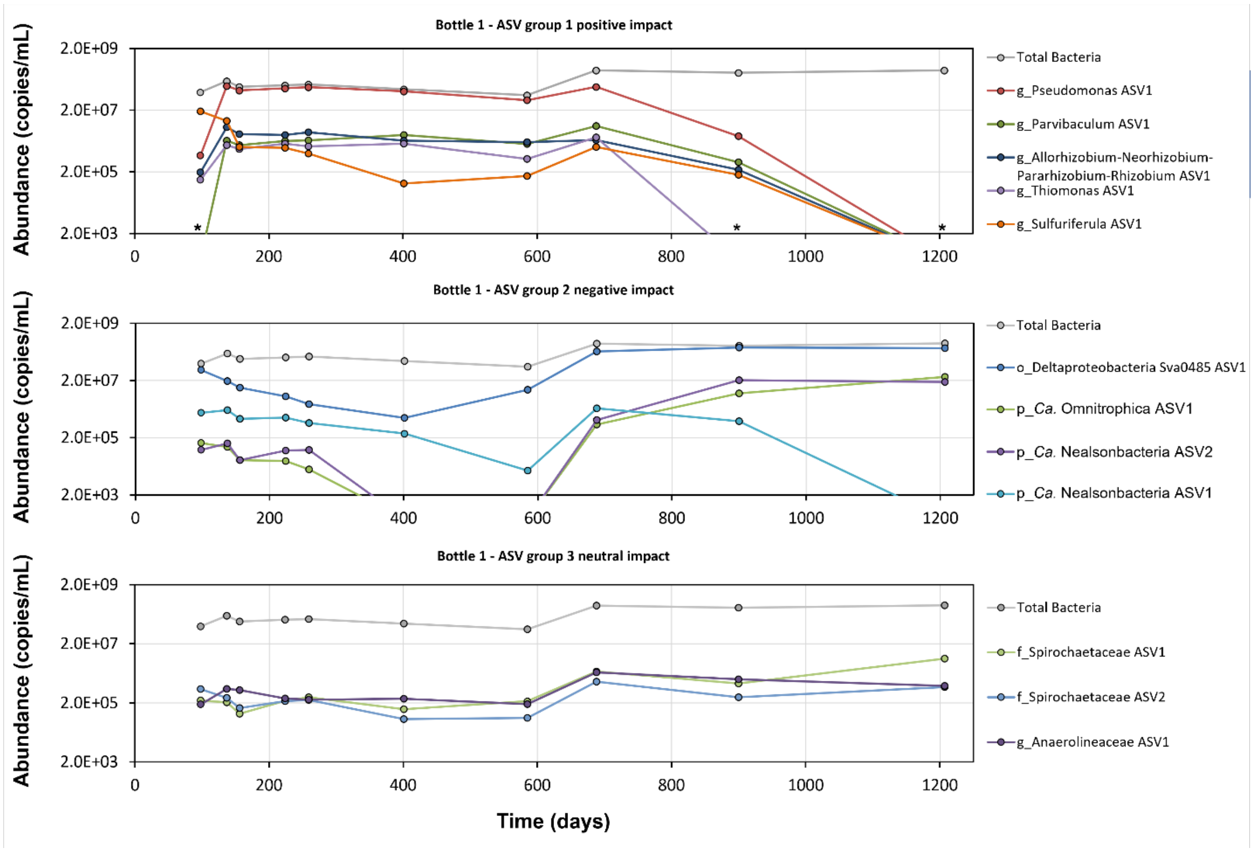


**Figure S6a**. Absolute abundance of specific bacterial ASVs in Bottle 1 expressed as 16S rRNA gene copy numbers per mL of culture. Among 22 most abundant bacterial ASVs (Table S5), the top panel summarizes ASVs which increased after exposure to oxygen (Group 1: positive impact of oxygen) and decreased when once methanogenic benzene degradation recovered in the culture. The center panel summarizes ASVs that decreased following exposure to oxygen but later recovered (Group 2: negative impact of oxygen). The bottom panel summarizes ASVs with negligible changes in concentration throughout the incubation period (Group 3: neutral). Abundances below quantifiable limits are designated by stars (*).

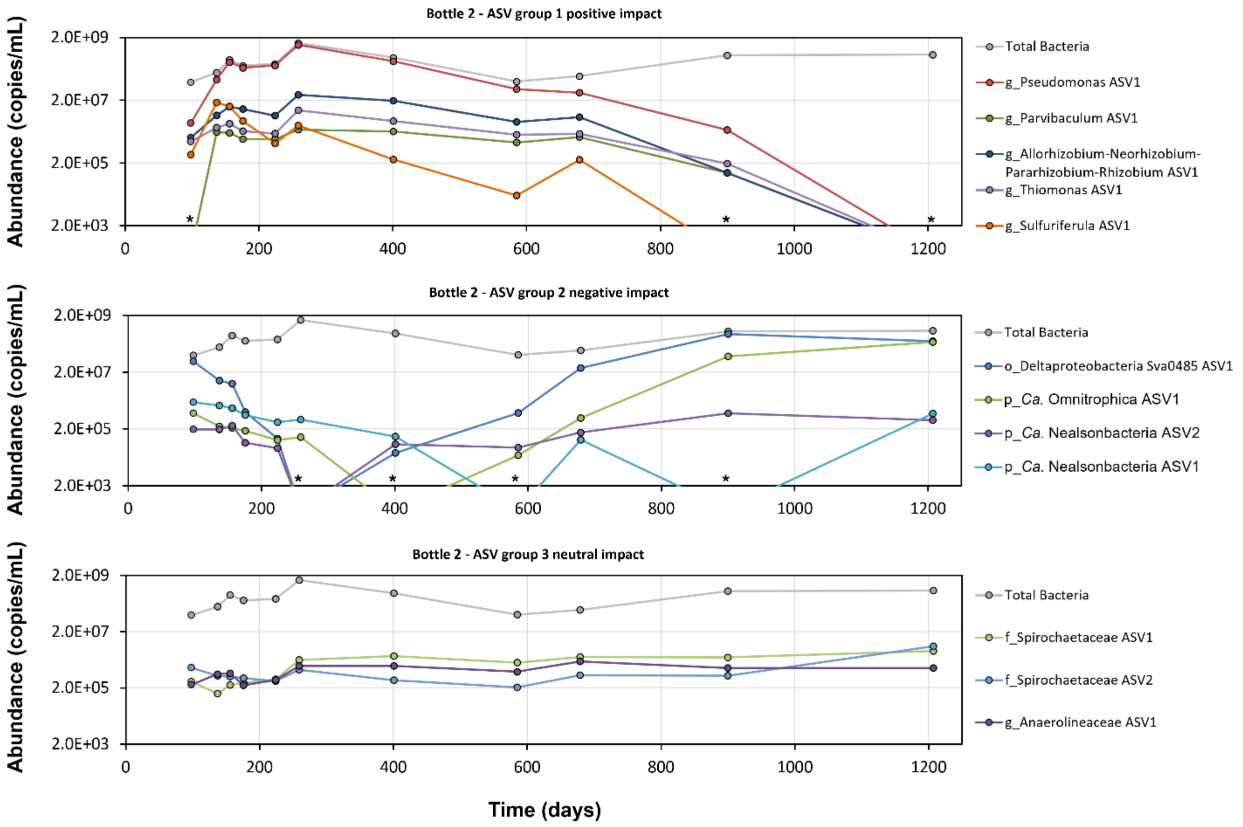


**Figure S6b**. Absolute abundance of specific bacterial ASVs in Bottle 2 expressed as 16S rRNA gene copy numbers per mL of culture. The three panels show Group 1, Group 2, and Group 3 ASVs as described for Figure S6a. Abundances below quantifiable limits are designated by stars (*).

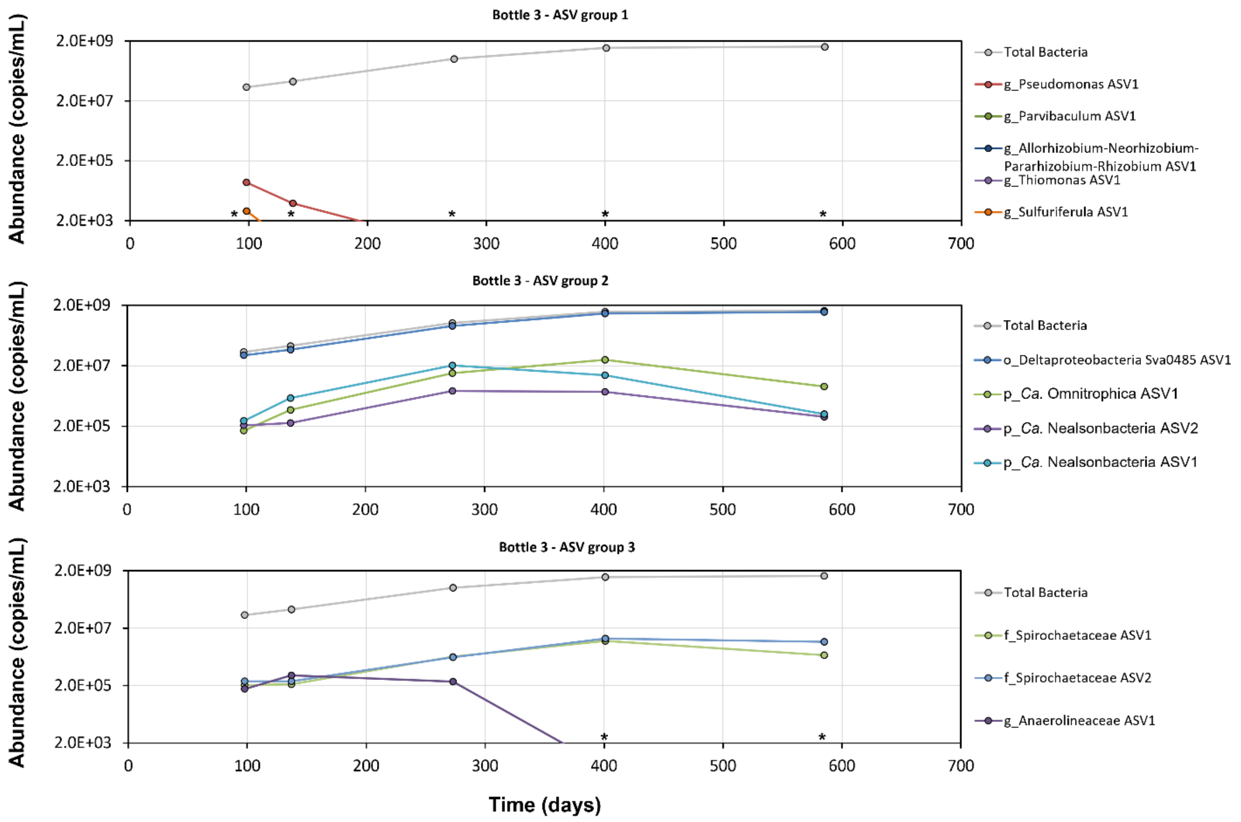


**Figure S6c**. Absolute abundance of specific bacterial ASVs in Bottle 3 expressed as 16S rRNA gene copy numbers per mL of culture. The three panels show Group 1, Group 2, and Group 3 ASVs as described for Figure S6a. Abundances below quantifiable limits are designated by stars (*).

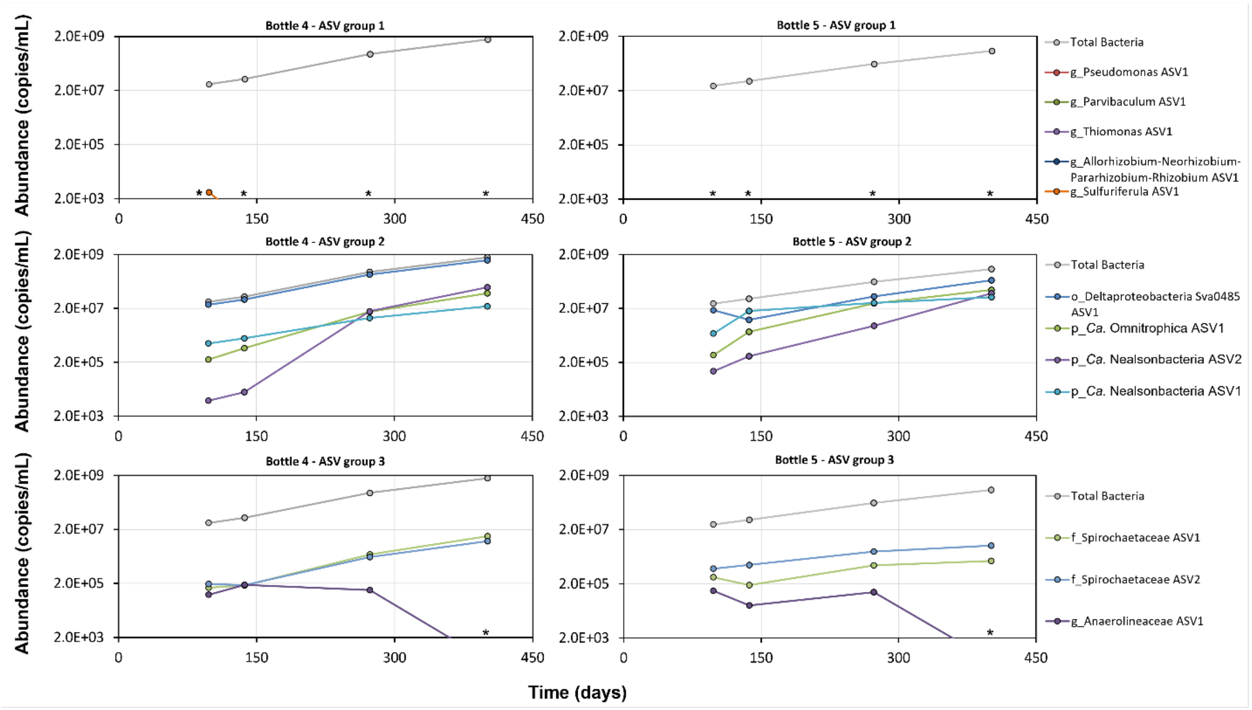


**Figure S6d**. Absolute abundance of specific <u>bacterial</u> ASVs in Bottles 4 and 5 expressed as 16S rRNA gene copy numbers per mL of culture. The three panels show Group 1, Group 2, and Group 3 ASVs as described for Figure S6a. Abundances below quantifiable limits are designated by stars (*).

**Figure 7 comprises a set 4 related figures (a,b,c,d) each for different bottles.**

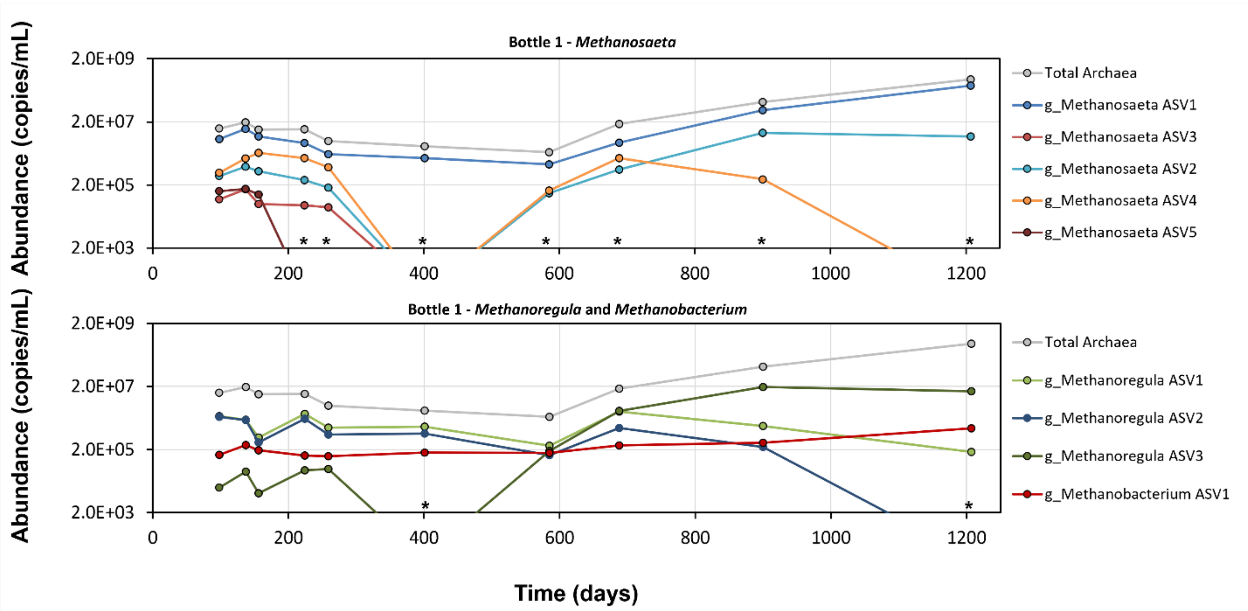


**Figure S7a**. Absolute abundance of specific archaeal ASVs in Bottle 1 expressed as 16S rRNA gene copy numbers per mL of culture. The top panel summarizes the five most abundant *Methanosaeta* ASVs (Table S6). The bottom panel summarizes the three most abundant *Methanoregula* ASVs and one *Methanobacterium* ASV (Table S6) Abundances below quantifiable limits are designated by stars (*).

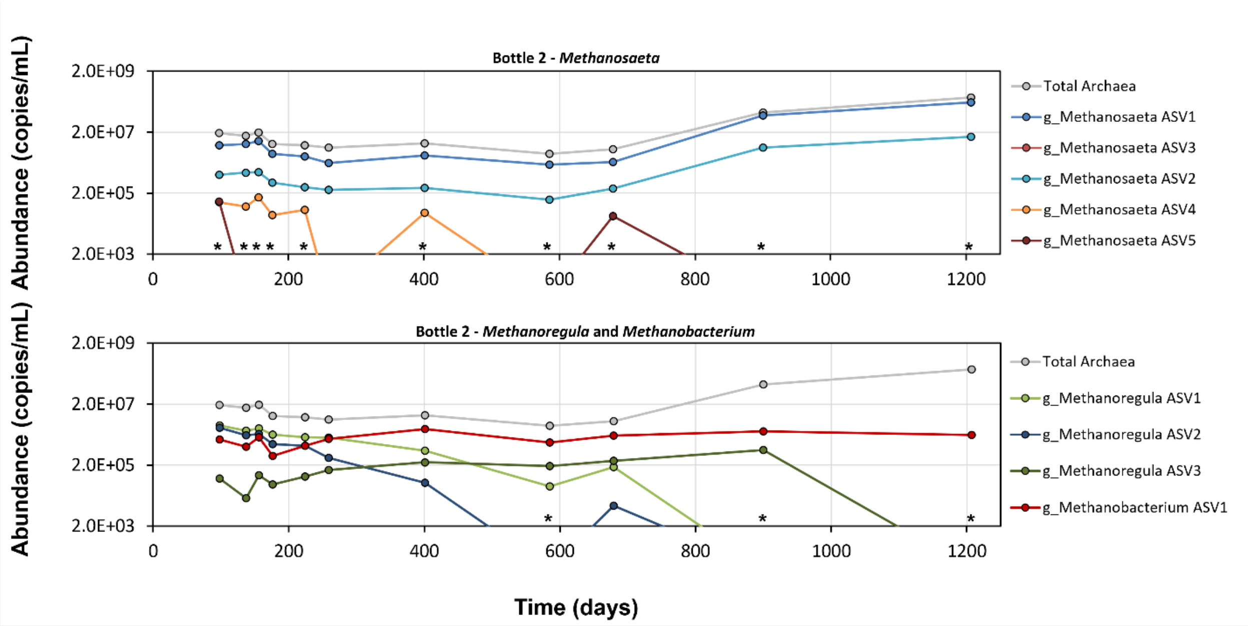


**Figure S7b**. Absolute abundance of specific archaeal ASVs in Bottle 2 expressed as 16S rRNA gene copy numbers per mL of culture. The two panels show the most abundant methanogens as described for Figure S6a. Abundances below quantifiable limits are designated by stars (*).

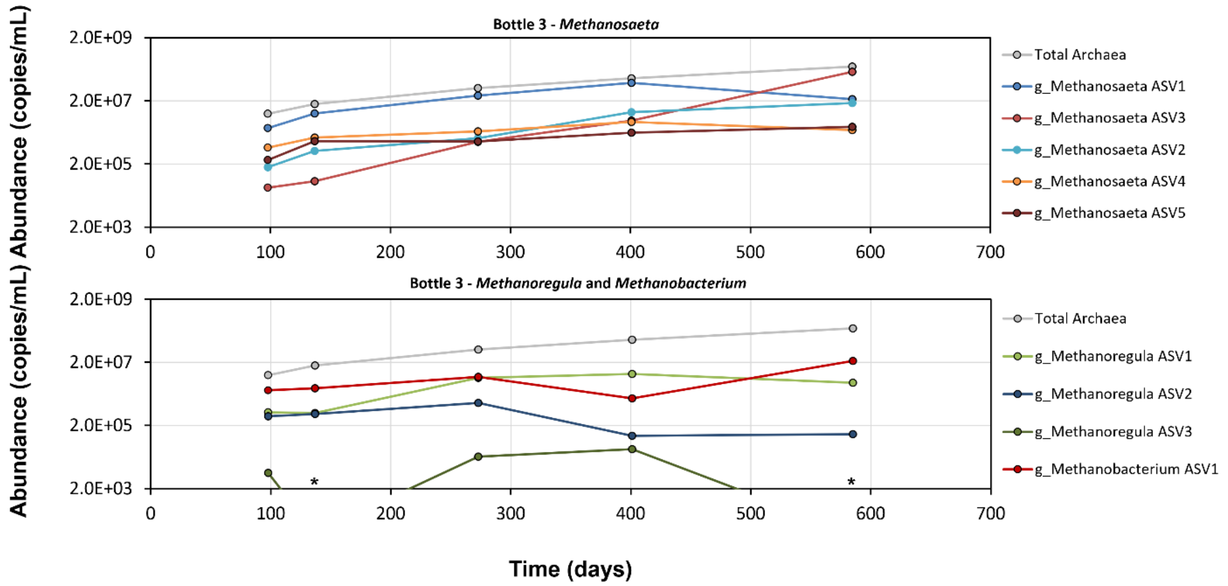


**Figure S7c**. Absolute abundance of specific archaeal ASVs in Bottle 3 expressed as 16S rRNA gene copy numbers per mL of culture. The two panels show the most abundant methanogens as described for Figure S6a. Abundances below quantifiable limits are designated by stars (*).

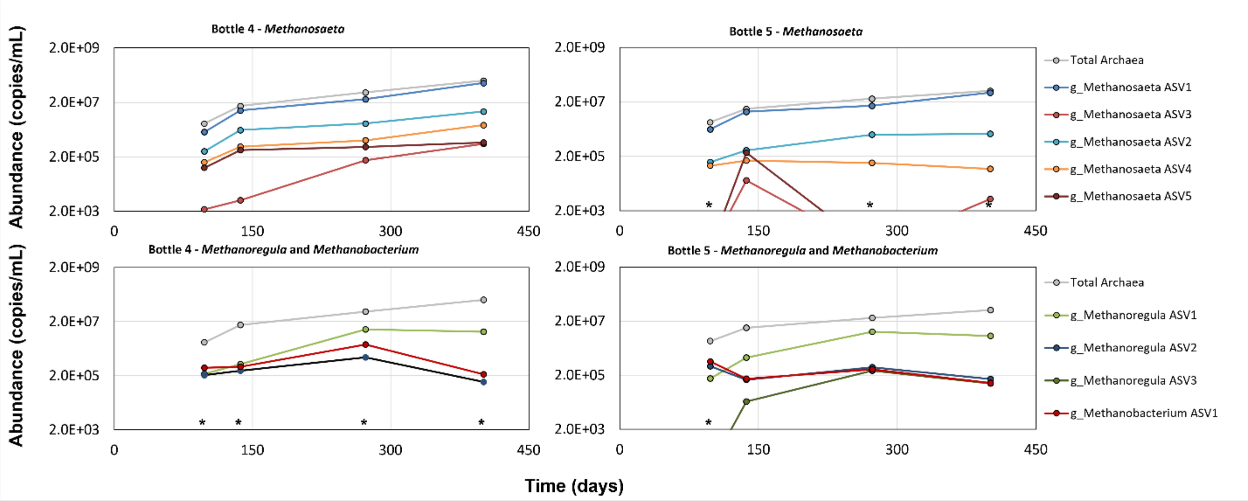


**Figure S7d**. Absolute abundance of specific archaeal ASVs in Bottle 4 and 5 expressed as 16S rRNA gene copy numbers per mL of culture. Panel a) summarizes the five most abundant *Methanosaeta* ASVs. Panel b) summarizes the three most abundant *Methanoregula* ASVs and one *Methanobacterium* ASV. Abundances below quantifiable limits are designated by stars (*).
